# Reconstitution of three-phase microtubule polymerisation dynamics

**DOI:** 10.1101/051672

**Authors:** Takashi Moriwaki, Gohta Goshima

## Abstract

Cytoplasmic microtubules (MTs) undergo growth, shrinkage, and pausing. However, how MT polymerisation cycles are produced and spatiotemporally regulated at a molecular level is unclear, as the entire cycle has not been recapitulated *in vitro* with defined components. In this study, we reconstituted dynamic MT plus end behaviour involving all three phases, by mixing tubulin with five *Drosophila* proteins, EB1, XMAP215^Msps^, Sentin, kinesin-13^Klp10A^, and CLASP^Mast/Orbit^. When singly mixed with tubulin, CLASP^Mast/Orbit^ strongly inhibited MT catastrophe and reduced the growth rate. However, in the presence of the other four factors, CLASP^Mast/Orbit^ acted as an inducer of pausing. The mitotic kinase Plk1^Polo^ modulated the activity of CLASP^Mast/Orbit^ and kinesin-13^Klp10A^, and increased the dynamic instability of MTs, reminiscent of mitotic cells. These results suggest that five conserved proteins constitute the core factors for creating dynamic MTs in cells, and that Plk1-dependent phosphorylation is a crucial event for switching from the interphase to mitotic mode.

## Introduction

Complementary to genetic analysis, *in vitro* reconstitution of intracellular processes, using defined factors, is a prerequisite to understand the process at a molecular level. This can determine if a given effect is directly mediated by each factor, and defines a minimal set of factors that can execute the process (Garner et al., 2007; Loisel et al., 1999; Shintomi et al., 2015; Yeeles et al., 2015). Dynamic microtubules (MTs) are composed of tubulin proteins, and play critical roles in various intracellular processes in eukaryotic cells, such as spindle formation and cell polarisation. Tubulin proteins, at particular concentrations, can autonomously and stochastically polymerise and depolymerise *in vitro*; this is referred to as MT dynamic instability (Horio and Hotani, 1986; Mitchison and Kirschner, 1984; Walker et al., 1988). In cells, free MT plus ends in the interphase cytoplasm repeatedly undergo three phases, growth, shrinkage, and pausing, in a stochastic order (Rogers et al., 2002; Shelden and Wadsworth, 1993). Multiple proteins control cellular MT dynamics, and depletion of these proteins results in alterations to MT dynamicity and consequently a change in the length of MT-based higher-order structures such as cilia or the mitotic spindle (Akhmanova and Steinmetz, 2008; Goshima and Scholey, 2010; Howard and Hyman, 2007; Hu et al., 2015). Most, if not all, of these proteins can individually modulate MT dynamics *in vitro.* Examples include XMAP215, which can increase MT growth rate, or kinesin-13, which can increase ‘catastrophe’ frequency (frequency of growth-to-shrinkage or pause-to-shrinkage transitions) (Howard and Hyman, 2007). When two or more factors are mixed with tubulin, the dynamic MT behaviour is better reproduced (Kinoshita et al., 2001; Li et al., 2012; Zanic et al., 2013). However, cellular MT dynamics, with all three phases, have not been successfully reproduced *in vitro.* Moreover, upon entry into mitosis, MTs become more dynamic, wherein MT growth rate and catastrophe frequency are increased and pausing is suppressed. However, the molecular basis of this cell cycle-dependent switch is also unclear.

The *Drosophila* S2 cell line is a promising model system for the reconstitution of MT plus end dynamics. Parameters of MT dynamics in interphase and mitosis have been obtained through live cell microscopy of fluorescent markers (Brittle and Ohkura, 2005; Li et al., 2011; Rogers et al., 2002; Sousa et al., 2007; Trogden and Rogers, 2015). Using this cell line, a handful of proteins required for dynamic regulation of MTs have been identified through multiple functional studies, including a large-scale functional genomics screen (Goshima et al., 2007; Hughes et al., 2008; Moutinho-Pereira et al., 2013). For instance, cells depleted of three conserved MT plus end tracking proteins Msps (XMAP215/ch-TOG orthologue), EB1 (a member of the EB protein family), and Sentin (a functional homologue of mammalian SLAIN2) similarly decreased MT dynamicity, wherein MT growth rate was reduced and pausing was drastically elevated (Brittle and Ohkura, 2005; Li et al., 2011; Rogers et al., 2002). To initiate *in vitro* reconstitution of MT plus end dynamics in S2 cells, we purified EB1, Sentin, and XMAP215^Msps^ proteins, mixed them with pig brain-derived tubulins, and assessed MT plus end polymerisation dynamics (Li et al., 2012). Compared to MTs generated solely with tubulin, or with one of these proteins, MTs formed in the presence of all three proteins had faster growth and more frequent catastrophe events, which is consistent with their depletion phenotype in cells (Brittle and Ohkura, 2005; Li et al., 2011; Li et al., 2012; Rogers et al., 2002). However, even with various concentrations and combinations of these proteins, an *in vitro* system that closely reproduces the *in vivo* dynamics has not yet been constructed; most critically, pausing and the ‘rescue’ event (shrinkage-to-growth or shrink-to-pause transitions) were observed with much lower frequency than in cells (Li et al., 2012).

We reasoned that a failure to reconstitute these three phases realistically might be due to missing factors in the *in vitro* reaction. In this study, we considered Klp10A (kinesin-13) and CLASP *(Drosophila* name is Mast or Orbit) as likely candidates for these missing factors, since RNAi depletion in S2 cells affected MT dynamics and mitotic spindle size (Goshima et al., 2005; Mennella et al., 2005; Sousa et al., 2007). Furthermore, orthologous proteins of other species were shown to modulate MT dynamics *in vitro* (Al-Bassam et al., 2010; Mennella et al., 2005). In addition, with recently established methodology (Widlund et al., 2012), we purified tubulin proteins from S2 cells and used them to replace pig tubulin proteins. In the reaction, *in vivo* MT dynamics were reproduced qualitatively, and to some extent, quantitatively, when five factors were mixed with S2 tubulin. Furthermore, phosphorylation by the mitotic kinase Polo (Plk1 in mammals) shifted MT dynamics to the mitotic mode.

## Results

### CLASP^Mast/Orbit^ suppresses MT growth and catastrophe

We first characterised the activity of S2 tubulin in the MT plus end polymerisation assay, in which dynamic MTs are generated from a stable MT seed made with GMPCPP-tubulin (Bieling et al., 2007) (Fig. 1A). As expected, S2-derived tubulins showed repeated growth-shrinkage cycles, as observed with pig tubulins, although each parameter was quantitatively different (Fig. S1A, S1C, D).

**Figure. 1.**
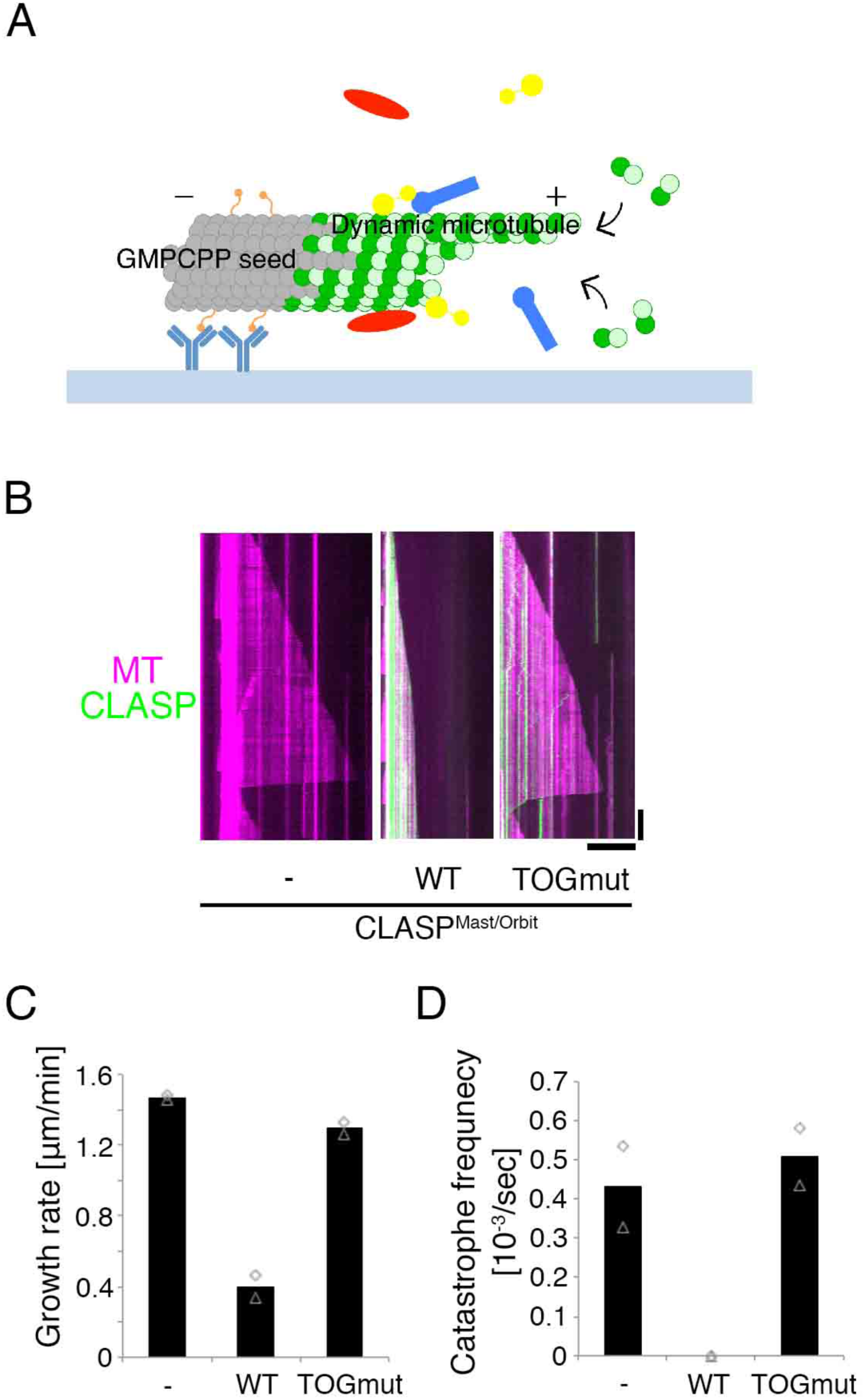
CLASP^Mast/Orbit^ slows growth and inhibits catastrophe *in vitro*. (**A**) Schematic of *in vitro* MT polymerisation assay used in this study. Green, tubulin; orange, biotinylation; light blue, anti-biotin antibody; red/yellow/blue, recombinant proteins. (**B**) CLASP ^Mast/0rbit^ slows MT growth and inhibits catastrophe. Kymographs show GFP-CLASP ^Mast/Orbit^ (wild-type and TOG domain mutant) and MT (10 μM S2 tubulin). (**C, D**)MT growth rate and catastrophe frequency in the presence of 45 nM CLASP ^Mast/0rbit^ (wild-type and TOG domain mutant) and 10 μm S2 tubulin. The mean values of each experiment are marked in grey, whereas the mean values of all the experiments are indicated by black graphs. Actual values are plotted in these graphs (n = 15–22 MTs). Horizontal bar, 10 μm; Vertical bar, 2 min.

Full-length kinesin-13^Klp10A^ was purified from *E*. *coli*,and it showed potent MT depolymerisation activity towards both ends of seed MTs (Fig. S1E), as expected (Mennella et al., 2005; Rogers et al., 2004).

GFP-tagged CLASP^Mast/Orbit^ was purified from insect Sf21 cells (Fig. S1A). Gel filtration chromatography indicated that the protein does not form aggregates (Fig. S1B). When mixed with S2 tubulin and MT seeds, GFP-CLASP^Mast/Orbit^ was localised along the seed and dynamic MTs, and a dramatic reduction in MT growth rate and catastrophe frequency was observed (Fig. 1B–D). Such activity was not observed, and MT association was reduced, when we used a mutant GFP-CLASP^Mast/Orbit^ in which four amino acid residues critical for MT/tubulin binding in the TOG2 and TOG3 domains were mutated (Al-Bassam and Chang, 2011; Al-Bassam et al., 2010; Leano et al., 2013) (Fig. 1B–D).Thus, the observed growth- and catastrophe-suppression activities were derived from CLASP^Mast/Orbit^, and not from contaminated proteins of Sf21 cells. In addition, the result indicated that the effect of GFP-CLASP^Mast/Orbit^ (on MT growth rate and catastrophe) requires its binding to tubulin and/or MTs. Previous RNAi experiments suggested that CLASP^Mast/Orbit^ induces MT pausing (Sousa et al., 2007; Trogden and Rogers, 2015), whereas purified fission yeast CLASP^Cls1^ protein was shown to have potent rescue (shrinkage-to-growth) activity (Al-Bassam et al.,2010). However, we could not assess if *Drosophila* CLASP^Mast/Orbit^ has these activities in this experiment, since MT shrinkage was never observed (Fig. 1D).

CLASP has an EB1-binding SxIP motif, and abundantly localises at the growing end of MTs in cells (Honnappa et al., 2009; Sousa et al., 2007). We reconstituted the plus-end-tracking behaviour of CLASP^Mast/Orbit^ by adding EB1 to the reaction (Fig. S1F). EB1 on its own slightly elevated MT growth rate and catastrophe frequency (Li et al., 2012). However, we could not detect any additional effects of CLASP^Mast/Orbit^ on growth rate; in addition, catastrophe could not be observed regardless of the presence or absence of EB1 (Fig. S1G). These results suggest that EB1-dependent plus-end-enrichment of CLASP^Mast/Orbit^ is not a prerequisite for the two CLASP^Mast/Orbit^-mediated activities towards plus ends.

### Reconstitution of three-phase MT dynamics with five factors

We next mixed all five factors in the reaction with S2-derived tubulin. Interestingly, we observed that MTs repeated cycles of shrinkage and growth, accompanied by frequent rescue and pausing; this behaviour had never been observed using just one to three factors (Fig. 2, Table 1). Each parameter value was within range of the corresponding cellular value, and the overall stochastic behaviour was reminiscent of interphase MT dynamics in S2 cells (Fig. S2).

**Figure. 2.**
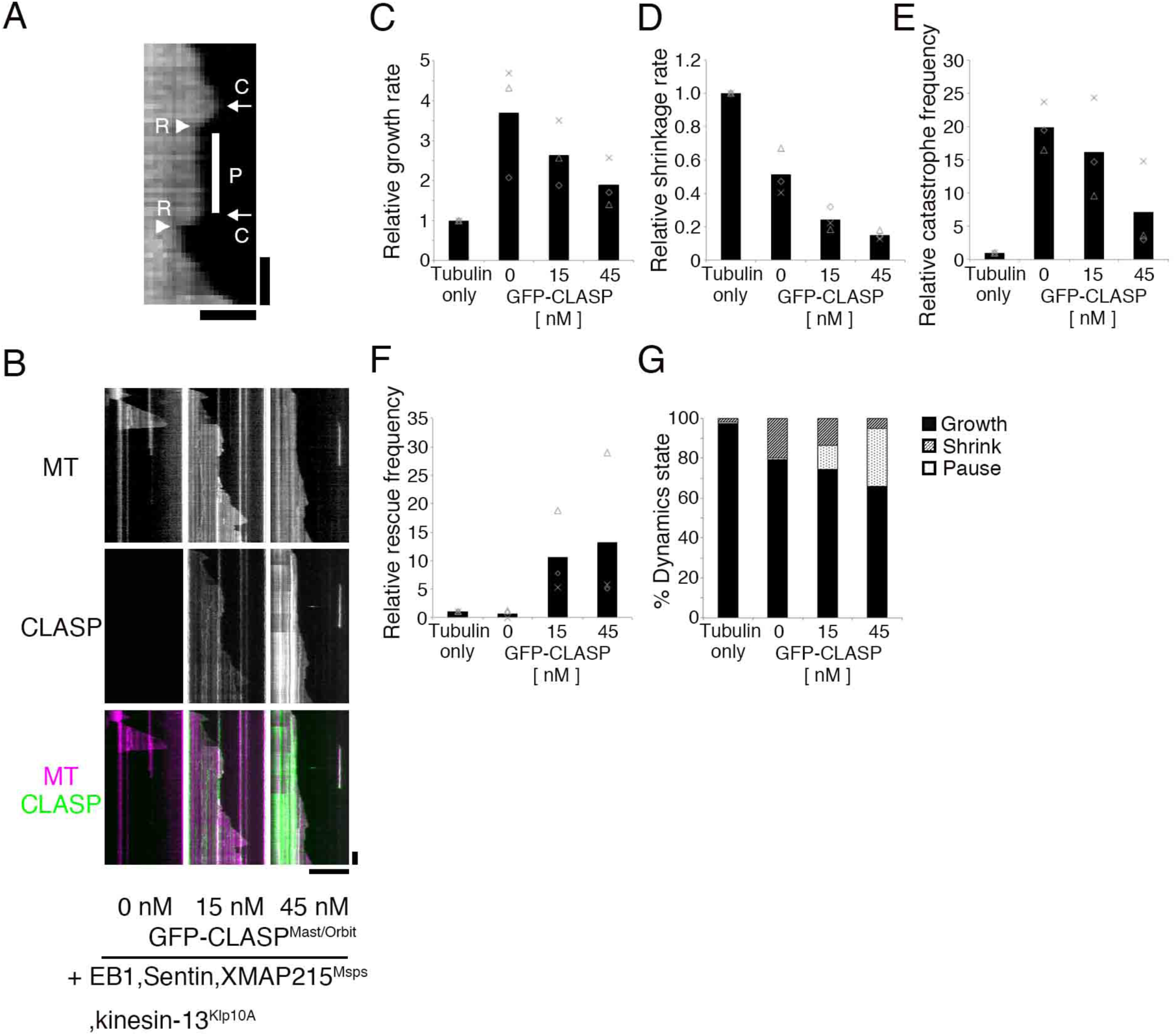
Reconstitution of dynamic MTs with five factors. (**A**) An example of pause (P) identified in the kymograph. Catastrophe (C) and rescue (R) events are also marked. Horizontal bar, 2 μm; Vertical bar 30 s. **(B-G)** Kymographs and kinetic parameters of MT polymerisation dynamics with 0–45 ηM GFP-CLASP^Mast/0rbit^ in the presence of 15 μM S2 tubulin, XMAP215^Msps^, EB1, Sentin and kinesin-13^Klp10A^. The mean values of each experiment are marked in grey, whereas the mean values of all the experiments are indicated by black graphs. Relative values are plotted in the graphs, whereas the actual values are presented in Table 1. Horizontal bar, 10 μm Vertical bar, 1 min.

**Table 1.** Kinetic parameters of *in vitro* reconstituted MT polymerisation dynamics with the addition of five factors (graphic presentation in Fig. 2)

| Condition | CLASP conc. | N | Growth rate | Shrinkage rate | Catastrophe frequency | Rescue frequency |
| --- | --- | --- | --- | --- | --- | --- |
| | (nM) | | ($\mu\text{m}/\text{min}$) | ($\mu\text{m}/\text{min}$) | ($\times 10^{-3}\text{s}^{-1}$) | ($\times 10^{-3}\text{s}^{-1}$) |
| Tubulin alone | 0 | 3 | 1.6 ± 0.1 (60) | 37.9 ± 8.2 (30) | 0.9 ± 0.2 (31) | 10.5 ± 6.1 (6) |
| +EB1/Sentin/Msps /Klp10A/CLASP | 0 | 3 | 5.8 ± 2.2 (88) | 19.0 ± 3.6 (86) | 17.7 ± 2.0 (86) | 5.3 ± 5.8 (7) |
| | 15 | 3 | 4.2 ± 1.3 (260) | 9.1 ± 2.5 (219) | 13.9 ± 3.6 (217) | 81.6 ± 11.2 (211) |
| | 45 | 3 | 3.0 ± 1.0 (166) | 5.6 ± 0.9 (88) | 5.7 ± 3.9 (86) | 86.1 ± 23.9 (87) |
| P-values (CLASP 45 nM vs. 0 nM) |  |  | 0.116 (n = 3)<br>0.002 (exp1)<br><10<sup>-4</sup> (exp2)<br><10<sup>-4</sup> (exp3) | 0.003 (n = 3)<br><10<sup>-4</sup> (exp1)<br><10<sup>-4</sup> (exp2)<br><10<sup>-4</sup> (exp3) | 0.009 (n = 3) | 0.005 (n = 3) |

Mean ± SD of three independent experiments (total number of events).

To validate the similarity between *in vitro* and *in vivo* dynamics, we omitted kinesin-13^Klp10A^, Sentin, or CLASP^Mast/Orbit^ from the reaction and compared the outcome with in vivo RNAi results. In S2 cells, kinesin-13^Klp10A^ knockdown decreases the catastrophe frequency (Mennella et al., 2005). Sentin knockdown inhibits the dynamic nature of MTs, specifically through increasing the pause duration and decreasing growth and shrinkage rates and catastrophe frequency (Li et al., 2011). CLASP^Mast/Orbit^ depletion in S2 cells increased the growth and shrinkage rates but reduced the pause duration (Sousa et al., 2007; Trogden and Rogers, 2015). Interestingly, we generally reproduced these tendencies in our *in vitro* assays through removal of kinesin-13^Klp10A^, Sentin, or CLASPMast/Orbit. Without kinesin-13^Klp10A^, catastrophe was dramatically suppressed and the imaging field was occupied by numerous MTs (Fig. 3A; only seven catastrophe events were observed during a 413-min growing phase). In the case of CLASP^Mast/Orbit^, all seven parameters were changed in a qualitatively consistent manner (Fig. 2B–F; “0 nM CLASP” corresponds to CLASP depletion; see also Fig. S3A–E for comparison to published *in vivo* results). Sentin depletion had similar impact; changes in six out of seven parameters were consistent *in vivo and in vitro* (Fig. 3B–G, Fig. S3F–J, Table S1).

**Figure. 3.**
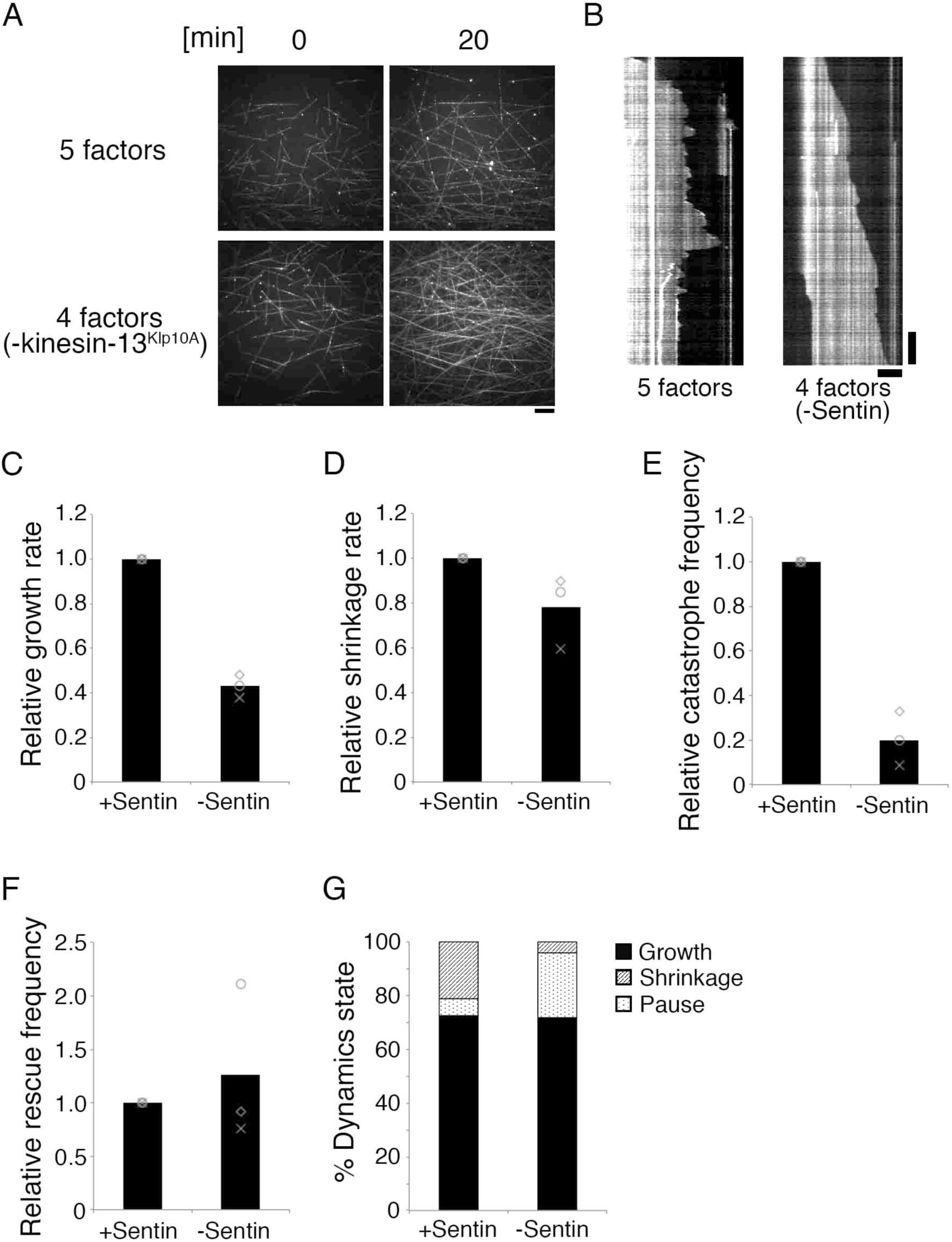
Effect of Sentin or kinesin-13^Klp10A^ removal from the MT polymerisation reaction. (**A**) MTs dramatically increased due to reduced catastrophe frequency in the absence of kinesin-13^Klp10A^ The other four factors were mixed with 15 μM S2 tubulin for 20 min. Bar, 10 μm. (**B-G**) MT polymerisation dynamics in the presence or absence of Sentin. The other four factors (EB1, XMAP215^Msps^, kinesin-13^Klp10A^, and 15 nM CLASP^Mast/Orbit^) were mixed with 15 μM S2 tubulin. Actual values are presented in Table S1.Horizontal bar, 5 μm; Vertical bar, 2 min.

### Lattice-bound CLASP^Mast/Orbit^ induces pausing in shrinking MTs

Since pausing was rarely observed in the absence of CLASP^Mast/Orbit^,we investigated whether GFP-CLASP^Mast/Orbit^ accumulation is correlated with pause induction. Since CLASP^Mast/Orbit^ was often abundantly localised to the growing end of MTs via EB1 binding, we focused our analysis on shrinking MTs. We first surveyed GFP-CLASP^Mast/Orbit^ signal distribution along the lattice of shrinking MTs (n = 21) and observed that 24% of the pixels contained a high GFP signal (Fig. 4A). This result indicated that if pausing is induced at random locations, the event has a 24% chance of occurring at GFP-accumulation sites. However, we identified 51% (n = 51) of shrinkage-to-pause events occurring at GFP-occupied sites (coloured red in Fig. 4B, C). Furthermore, in 9 out of the 25 events, in which pausing was induced at sites with low GFP-CLASP^Mast/Orbit^ abundance, MTs already had strong GFP signals during shrinking (coloured green in Fig. 4B, C). Thus, GFP-CLASP^Mast/Orbit^ accumulation and pause induction were well correlated, albeit not perfectly. To investigate whether CLASP^Mast/Orbit^ on the lattice recruits other factors, we mixed MT seeds with TagRFP-CLASP^Mast/Orbit^ and GFP-Sentin; Sentin is proposed to act as a bridge between CLASP^Mast/Orbit^ and XMAP215^Msps^ (Trogden and Rogers, 2015) but does not localise to the MT lattice on its own (Li et al., 2012). We did not observe lattice localisation of the GFP-Sentin signals in the presence of TagRFP-CLASP^Mast/Orbit^, suggesting that pre-recruitment of the Sentin-XMAP215^Msps^ complex is not required for CLASP^Mast/Orbit^ to induce pausing (Fig. S1H).

**Figure. 4.**
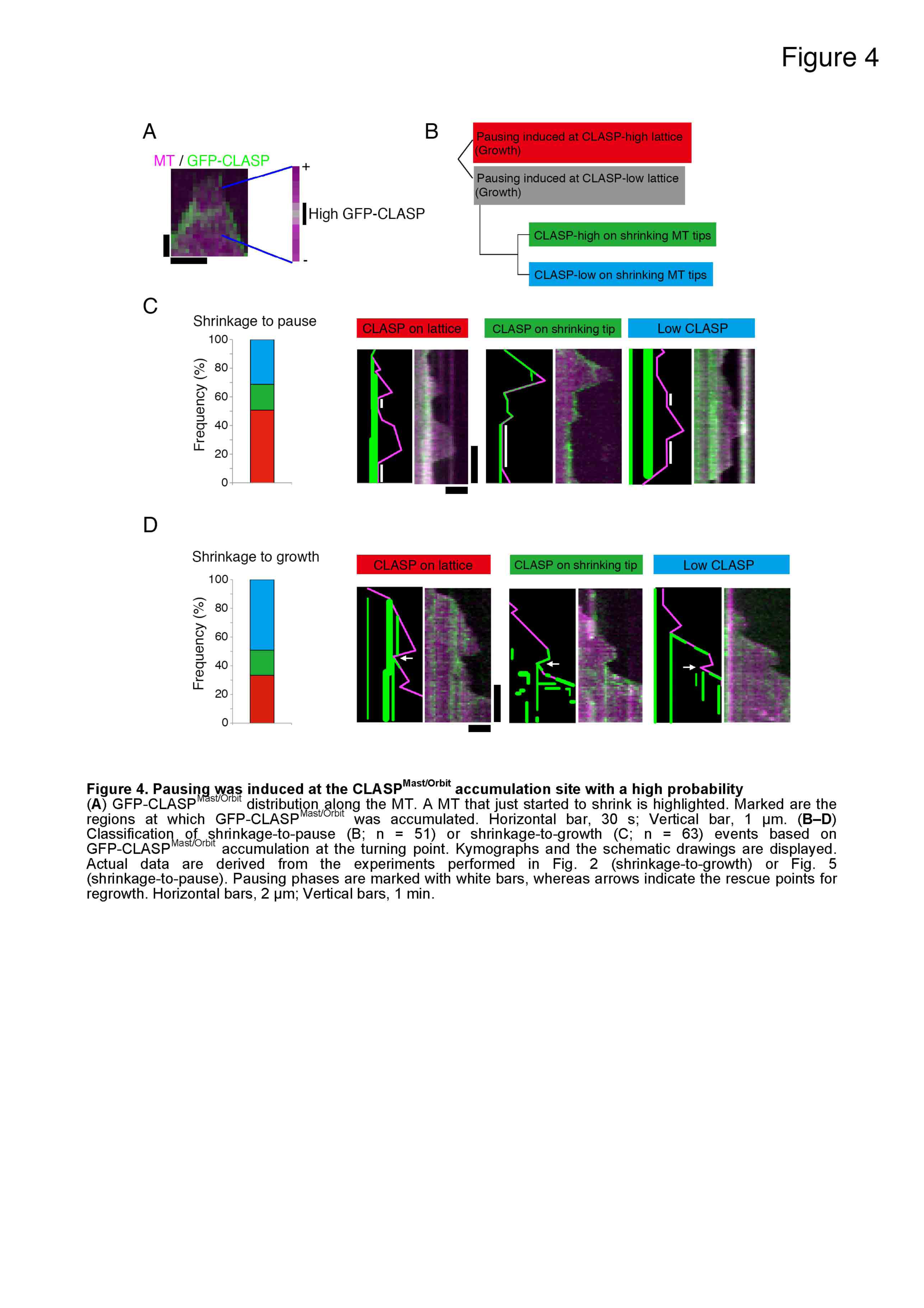
Pausing was induced at the CLASP^Mast/0rbit^ accumulation site with a high probability. **(A)** GFP-CLASP^Mast/0rbit^ distribution along the MT. A MT that just started to shrink is highlighted. Marked are the regions at which GFP-CLASP^Mast/Orbit^ was accumulated. Horizontal bar, 30 s; Vertical bar, 1 μm. (**B-D**) Classification of shrinkage-to-pause (B; n = 51) or shrinkage-to-growth (C; n = 63) events based on GFP-CLASP^Mast/Orbit^accumulation at the turning point. Kymographs and the schematic drawings are displayed.Actual data are derived from the experiments performed in Fig. 2 (shrinkage-to-growth) or Fig. 5 (shrinkage-to-pause). Pausing phases are marked with white bars, whereas arrows indicate the rescue points for regrowth. Horizontal bars, 2 μm; Vertical bars, 1 min.

Similar analysis was performed for shrinkage-to-growth events (Fig. 4B, D). In this case, the correlation was weaker than for the pause, but was still higher than expected from random occurrence (33% events occurred at GFP accumulation site [n = 63], which occupied 15% of the pixels [n = 31]). This result indicates that small amounts of CLASP^Mast/Orbit^ can also trigger the shrinkage-to-growth rescue, which differs from the results of fission yeast CLASP^Cls1^, wherein enrichment was perfectly correlated with the rescue event (Al-Bassam et al., 2010). We could not rule out the possibility that our microscope was not sensitive enough to detect moderately accumulated GFP-CLASP^Mast/Orbit^. Regardless, the results suggest that fly CLASP is a potent pause/rescue-inducing factor in the presence of four other factors.

### Increase in MT dynamicity by Plk1^Polo^ kinase

MT dynamics are subjected to mitotic regulation. During mitosis, the rate of astral MT growth and the frequency of catastrophe events increase, whereas the frequency of rescue events decreases (Belmont et al., 1990; Rusan et al., 2001). It was shown that the addition of active Cdk1 to the interphase frog extracts triggers the interphase-mitosis transition of MT dynamics (Verde et al., 1990). However, the event downstream of Cdk1 remains unclear. We first hypothesised that Cdk1 might directly phosphorylate one or more of the five factors, or tubulin itself, to modulate MT dynamics. Autoradiography experiments indicated that kinesin-13^Klp10A^ and XMAP215^Msps^ are phosphorylated by the recombinant Cdk1-cyclin B complex (Fig. S4A). Next, we incubated the five proteins and tubulin with Cdk1-cyclin B, followed by a MT polymerisation assay. Here, kinetic parameters were not significantly or reproducibly changed in four experiments comparing the effect of Cdk1 (Fig. S4B–F, Table S2). These results were not consistent with the idea that Cdk1 phosphorylation directly regulates the activities of these six proteins, although they did not entirely disprove it.

We considered it likely that a factor other than Cdk1 is the direct contributor to the increase in MT dynamicity during mitosis. Polo-like kinase 1 (Plk1) consists of a complex regulatory circuit during mitotic entry (Lindqvist et al., 2009; Zitouni et al., 2014). We therefore next hypothesised that Plk1 may be responsible for the change in dynamics. Interestingly, incubation of five factors and tubulin with recombinant Plk1^Polo^ affected kinetic parameters in a reproducible manner; growth rate and catastrophe increased, whereas rescue and pausing were reduced, making MTs more dynamic (Fig. 5B, C, Table S3). Phos-tag and autoradiography analyses showed that kinesin-13^Klp10A^, Sentin, and CLASP^Mast/Orbit^ were phosphorylated by Plk1^Polo^, whereas no gel shift was observed after Phos-tag analysis for EB1, XMAP215^Msps^ or S2 tubulin (Fig. 5A). When GFP-CLASP^Mast/Orbit^ was mixed with tubulin, we observed a reduction in MT lattice association in the presence of Plk1^Polo^ (Fig. 5D, E). This result was consistent with the increase of MT dynamicity conferred by Plk1^Polo^, as lattice-associated CLASP^Mast/Orbit^ frequently induced pausing (Fig. 4C). This effect was at least in part phosphorylation-dependent, since kinase-inactive (KD) Plk1^Polo^, purified in an identical manner to wild-type, showed a milder effect (Fig. 5D, 5). Conversely, when active Plk1^Polo^ was mixed with the other four factors and tubulin, we observed rapid depolymerisation of MT seeds (Fig. 5F, G). The seed depolymerisation suggested that kinesin-13^Klp10A^ is activated by Plk1^Polo^. Thus, Plk1^Polo^-mediated phosphorylation affected the activities of CLASP^Mast/Orbit^ and kinesin-13^Klp10A^. Whether Sentin, which also has catastrophe-inducing activity (Li et al., 2012), is a critical substrate of Plk1^Polo^ remains to be determined.

**Figure. 5.**
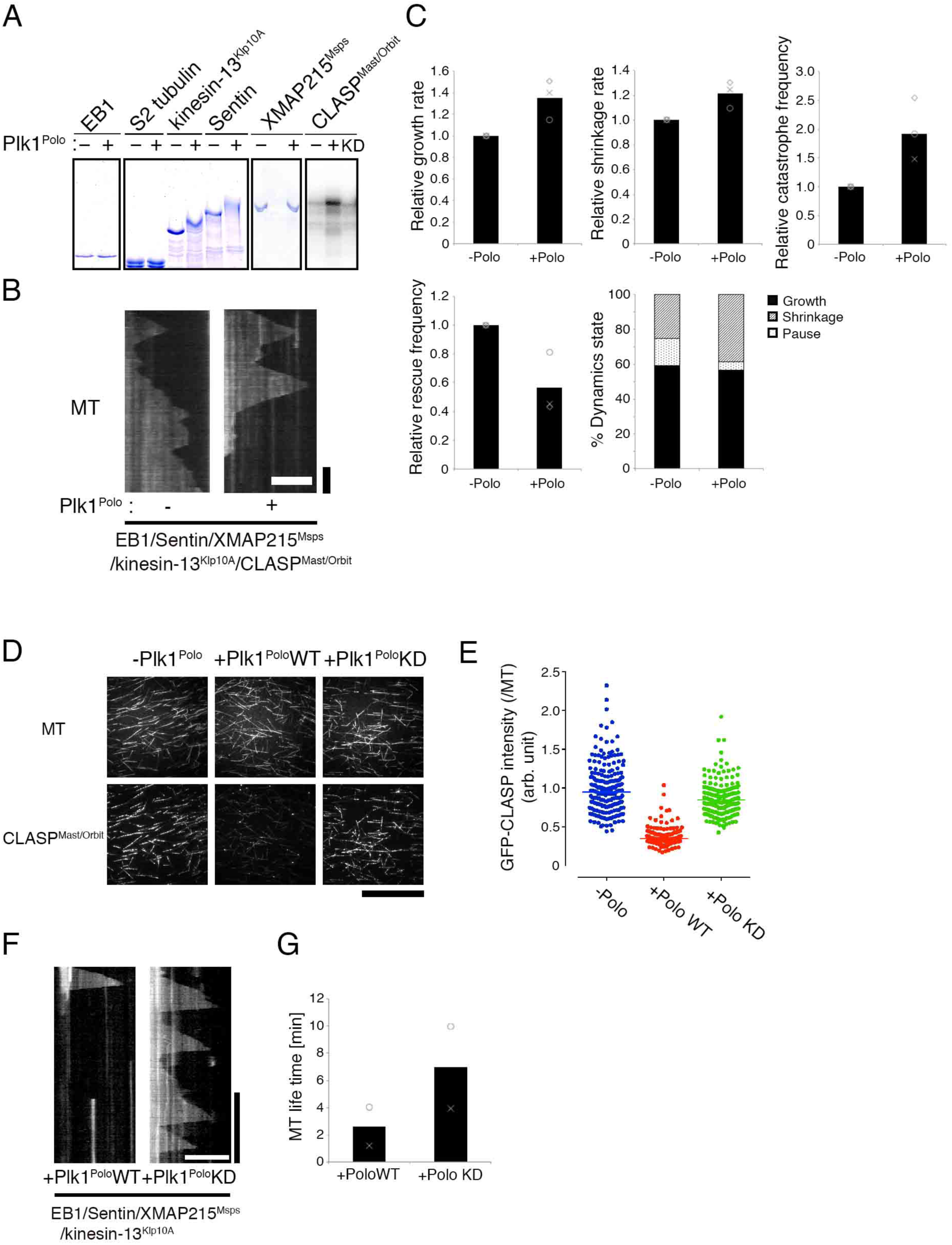
Phosphorylation by Rlk1^Polo^ elevated MT dynamicity *in vitro*. (**A**) Autoradioaraphy (rightmost panel) and Phos-tag gel electrophoresis (others) showed phosphorylation of CLASP^Mast/Orbit^, kinesin-13^Klp10A^, and Sentin by Plk1^Polo^. (**B,C**) Kymographs and kinetic parameters showing that treatment of five MT dynamic-modulating factors with Plk1^Polo^ makes MTs more dynamic (10 μM S2 tubulin; 45 nM CLASP^Mast/0rbit^). In the graph, the mean and relative values of each experiment are plotted. Actual values are presented in Table S3. Horizontal bar, 5 μm; Vertical bar, 1 min. (**D, E**) Association of GFP-CLASP^Mast/0rbit^ (5 nM) with the MT lattice (10 μM S2 tubulin) was drastically reduced by incubation with wild-type Plk1^Polo^ (100 nM; p < 0.0001, Mann-Whitney U-test), and more modestly with a kinase-dead (KD) mutant. Bar, 50 μm.(**F,G**) MT depolymerisation activity of kinesin-13^Klp10A^ was enhanced by incubation with wild-type Plk1^Polo^°. Four factors (EB1, XMAP215^Msps^, Sentin, and 10 nM kinesin-13^Klp10A^) and 15 μM S2 tubulin were included in this reaction and seed depolymerisation was assessed after generating the kymograph. Note that the lifetime for wiid-type Plk1^Polo^ is likely overestimated, since some seeds had been already depolymerised when the imaging was started. Horizontal bar, 10 μm; Vertical bar, 5 min.

**Table S3.** Kinetic parameters of *in vitro* reconstituted MT polymerisation dynamics with or without Plk1^Polo^(graphic presentation in Fig. 5)

| Condition | N | Growth rate | Shrinkage rate | Catastrophe frequency | Rescue frequency |
| --- | --- | --- | --- | --- | --- |
| | | ($\mu\text{m}/\text{min}$) | ($\mu\text{m}/\text{min}$) | ($\times 10^{-3}\text{s}^{-1}$) | ($\times 10^{-3}\text{s}^{-1}$) |
| -Polo | 3 | 3.1 ± 0.2 (314) | 5.5 ± 0.1 (289) | 19.6 ± 5.9 (241) | 45.8 ± 9.3 (290) |
| +Polo | 3 | 4.2 ± 0.5 (191) | 6.7 ± 0.6 (190) | 36.0 ± 8.0 (86) | 27.1 ± 15.5 (186) |
| P-values |  | 0.032 (n = 3)<br>0.001 (exp1)<br><10<sup>-4</sup> (exp2)<br>0.093 (exp3) | 0.027 (n = 3)<br>0.016 (exp1)<br>0.002 (exp2)<br>0.127 (exp3) | 0.046 (n = 3) | 0.147 (n = 3) |

Mean ± SD of three independent experiments (total number of events).

To investigate the involvement of Plk1^Polo^ in MT dynamics *in vivo*, we measured growth velocity of mitotic astral MTs in S2 cells in the presence or absence of the Plk1^Polo^ inhibitor BI2536. S2 cells have heterogeneity in centrosome numbers, which might alter the parameters associated with MT dynamics of each aster. Therefore, we measured astral MT dynamics in the monopolar spindle that was induced by RNAi knockdown of the kinesin-5 motor Klp61F (Fig. 6A). Interestingly, mitotic EB1-GFP velocity was decreased by 40% following BI2536 treatment, which matched the velocity in interphase (Fig. 6B). The decrease was unlikely induced by off-target effect of BI2536 on tubulin, since the MT growth rate during interphase was unaffected by this treatment. We next used the same dataset to compare growth time. Since pausing is a rare event during S2 mitosis (Li et al., 2011), we considered that disappearance of the EB1-GFP signals in the mitotic aster mostly corresponded to catastrophe. Duration of EB1-GFP tracking was 56% longer in the presence of BI2536 and this was similar to interphase values, suggesting that Plk1^Polo^ elevated catastrophe frequency in mitotic cells (Fig. 6C).

**Figure. 6.**
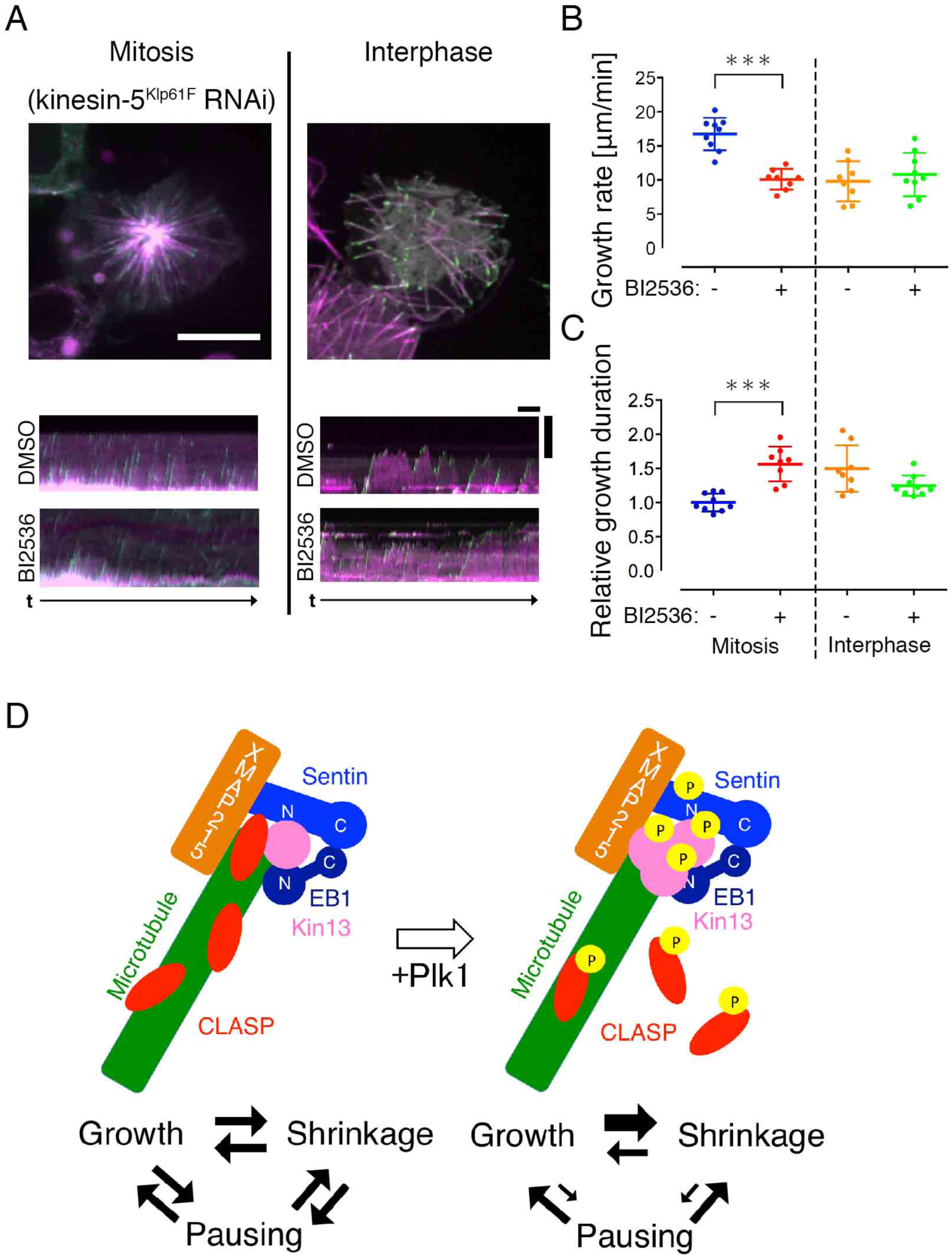
Plk1^Polo^ inhibition dampens MT dynamicity *in vivo*. (**A-C**) Plk1^Polo^inhibition decreased MT growth rate and catastrophe frequency in mitosis but not in interphase. Mitotic astral MTs in the monopolar spindle induced by Klp61F (kinesin-5) RNAi were analysed (100–200 comets per cell). Each dot in the graph represents the mean value obtained from a cell. Asterisks indicate statistically significant differences (p < 0.0001; Turkey’s multiple comparisons test). Bar for cell images, 10 μm; Horizontal bar for kymographs, 1 min; Vertical bar for kymographs, 5 μm. (**D**)Reconstitution of regulated microtubule polymerisation dynamics was achieved in this study. Three-phase polymerisation cycles were observed when five factors were mixed with tubulin. Addition of the active Plk1 kinase increased MT dynamicity, wherein pausing became rare.

## Discussion

This study aimed to reproduce cellular MT polymerisation dynamics *in vitro* more faithfully than previous studies. Towards this goal, we expanded the materials for reconstitution to seven purified proteins (including tubulin and a kinase). To the best of our knowledge, this presents the most complex reconstitution of MT polymerisation dynamics. Our *in vitro* experiments, together with previous loss-of-function analyses *in vivo*, suggest that five proteins, XMAP215^Msps^, EB1, Sentin, kinesin-13^Klp10A^, and CLASP^Mast/Orbit^, constitute the core machinery that regulates MT dynamics in the S2 cytoplasm (Fig. 6D, left). In the presence of these five factors, typical MT behaviour was reproduced, in which growth, shrinkage, and pausing phases repeat in a stochastic order. The change in dynamics in the absence of CLASP^Mast/Orbit^, Sentin, or kinesin-13^Klp10A^ supports this model. Omission of each factor *in vitro* resulted in MT dynamics that deviated from that of untreated S2 cytoplasm; dynamics became more similar to that observed upon RNAi-depletion of corresponding factors. Moreover, phosphorylation by the mitotic kinase Plk1^Polo^ resulted in more dynamic MTs, reminiscent of what occurs during interphase-to-mitosis transitions (Fig. 6D, right).

### CLASP induces pausing

A previous study using purified yeast CLASP^Cls1^ identified rescue (shrinkage-to-growth) and catastrophe-inhibiting activities, but not pausing or growth-inhibiting activity (Al-Bassam et al., 2010). An intriguing observation in the current study was the emergence of pause when CLASP^Mast/Orbit^ was added with the four other factors. This illustrates the value of multi-factor reconstitution and suggests that an equilibrium of CLASP^Mast/Orbit^ with MT polymerase and depolymerase produces such a state. It is possible that such activity was conferred by physical interaction with other factors; for instance, it was recently shown that XMAP215^Msps^ interacts with CLASP^Mast/Orbit^ via Sentin in S2 cells (Trogden and Rogers, 2015). In S2 cells, RNAi depletion of EB1, Sentin, or XMAP215^Msps^ increases the pause duration in interphase, and this phenotype could not be readily explained (Brittle and Ohkura, 2005; Li et al., 2011; Rogers et al., 2002). On the basis of current *in vitro* results, this phenotype may be interpreted that when either EB1, Sentin or XMAP215^Msps^ is depleted, CLASP^Mast/Orbit^ dominates at the plus end, and induces pausing together with the other three factors.

### Plk1 elevates MT dynamicity

Another finding in this study was the increase in MT dynamicity by Plk1^Polo^ kinase. The involvement of Plk1^Polo^ is not inconsistent with the classical analysis in which Cdk1 triggers elevation of MT dynamicity in frog egg extracts (Verde et al., 1990). It has been shown that Cdk1 and Plk1 comprise a complex signalling network, and the addition of Cdk1 to egg extracts may well activate the Plk1 kinase. Consistent with the *in vitro* data, we observed reversion of MT dynamics, from mitotic to interphase mode, by treating prometaphase cells with a Plk1^Polo^ inhibitor. An *in vitro* kinase assay showed that Sentin, kinesin-13^Klp10A^, and CLASP^Mast/Orbit^ are the substrates of Plk1^Polo^. The appearance of multiple bands in the electrophoresis gel indicates that multiple residues are potential Plk1^Polo^ phosphorylation sites. Determining the substrate residues that are phosphorylated to influence MT dynamics *in vivo* is an important future research topic. However, our results do not exclude the possibility that Plk1^Polo^ phosphorylation of other proteins also contributes to changes in MT dynamics in cells.

### Comparison between *in vitro* and *in vivo* data

While we believe we have achieved hitherto the most *in vivo*-relevant reconstitution, we admit that the study is still not a precise reproduction of S2 MT dynamics. First, other factors may also contribute to MT dynamics in cells. S2 cells express additional factors that can modulate MT dynamicity *in vitro,* such as D-TPX2 and two other kinesin-13 family members (Klp59C and Klp59D); however, RNAi-depletion of these factors results in a phenotype that is not as severe as that observed after ablation of the five factors used in this study (Goshima, 2011; Goshima and Vale, 2003; Mennella et al., 2005). Second, our data on MT dynamics are basically limited to free MTs in the cytoplasm. Other types of MTs, such as cortical/peripheral MTs, are often bound to other protein/membranous structures and show different dynamic kinetics (Trogden and Rogers, 2015), and additional factors are likely necessary for reconstitution. For instance, human CLASP depletion decreased catastrophe as well as rescue at the cell cortex, which is not entirely consistent with the consequence we observed here *in vitro* (Mimori-Kiyosue et al., 2005); GSK3 kinase is known to phosphorylate CLASP and thereby regulate peripheral MT dynamics (Kumar et al., 2009; Trogden and Rogers, 2015). Kinetochore MTs in metaphase are also bound to many other proteins. However, a key feature of kinetochore MT dynamics might have been partially reproduced with one of our assay conditions. In cells, CLASP^Mast/Orbit^ is most prevalent at the outer kinetochore region during metaphase, and is required for kinetochore MT polymerisation (Maiato et al., 2005). The slow but persistent growth of kinetochore MTs has resemblance to that observed when a high concentration of CLASP^Mast/Orbit^ was mixed with four other factors; MTs grew slower and rarely shrank even in the presence of catastrophe-promoting factors (Fig. 2, 45 nM CLASP^Mast/Orbit^). Thus, although multiple other MT-regulating proteins are present at the outer kinetochore, it is tempting to speculate that accumulation of CLASP^Mast/Orbit^ is a key event that distinguishes kinetochore MTs from highly dynamic astral MTs.

Finally, another difference between our system and the *in vivo* situation involves the environment and protein concentrations. We used a standard non-viscous buffer in our assay, which would be different from the cytoplasm that is highly viscous, containing a number of other molecules. Furthermore, our rough estimate indicated that cellular concentrations of CLASP^Mast/Orbit^ and kinesin-13^Klp10A^ were 45–80-fold higher than what we used *in vitro*, a level currently unachievable by our purification method (Fig. S5). One possible explanation for this discrepancy is that cellular proteins are regulated spatially or post-translationally and actual proteins that can target MTs are limited. Alternatively, the antagonistic activity of CLASP^Mast/Orbit^ and kinesin-13^Klp10A^ on catastrophe suggests that the molecular ratio of the two proteins might be critical, rather than absolute abundance of each protein. In support of this idea, the ratio of CLASP^Mast/Orbit^ and kinesin-13^Klp10A^ was comparable between *in vitro* (1:6.7) and *in vivo* (1:3.7) conditions. Regardless, our reconstitution admittedly is qualitative, and quantitative comparison with *in vivo* kinetic parameters remains difficult. Converting the kinetic parameters and protein concentrations to *in vivo* values remains a key challenge in the reconstitution of MT dynamics and perhaps also for other complex cellular events (Shintomi et al., 2015).

### Towards reconstitution of mammalian MT dynamics

The data obtained in this study might be largely applicable to mammalian cells. EB1, XMAP215^Msps^, kinesin-13^Klp10A^, CLASP^Mast/Orbit^, and Plk1^Polo^ have clear orthologues in mammals. Although not conserved in amino acid sequences, Sentin is likely a functional homologue of mammalian SLAIN2, because both proteins bind to EB1 and XMAP215^Msps^, bridging these two molecules for synergistic MT growth acceleration (Li et al., 2011; van der Vaart et al., 2011). Furthermore, Plk1-dependent activation of kinesin-13 has been demonstrated *in vitro* (Jang et al., 2009; Zhang et al., 2011). Since multiple paralogous/homologous genes have been identified for CLASP (1 and 2), EB (1, 2, 3), SLAIN (1 and 2), and kinesin-13 (Kif2a, Kif2b and MCAK), mammalian cells might have further layers of complexity compared to *Drosophila.* In fact, it was recently shown that human CLASP1 and an isoform of CLASP2 have common but also distinct activities (Yu et al., 2016). Regardless, a reasonable first attempt towards reconstitution of mammalian MT dynamics would be to purify and mix homologues of the five proteins identified in this study.

## Materials and methods

### Protein purification

S2 tubulin was purified by using a previously described method (Widlund et al., 2012). Cell culture (700 mL) was lysed with 7 mL MRB80 buffer (80 mM K-PIPES [pH 6.8], 4 mM MgCl_2_, and 1 mM EGTA) supplemented with 5ul/125U Benzonase (Novagen), followed by affinity purification with 1 mL TOG1 column purification (TOG1-encoding plasmid was a gift from Dr. Per Widlund). EB1 was purified using *Escherichia coli* as described previously (Li et al., 2011; Li et al., 2012). In brief, GST-EB1 was purified with glutathione sepharose beads, followed by proteolytic cleavage of the GST tag and gel filtration chromatography. His-kinesin-13^Klp10A^ was purified using the standard Ni-NTA purification with the buffer identical to that of EB1, except that 1 mM ATP was supplied. His-Sentin, His-GFP-Sentin, and XMAP215^Msps^-HAHis were purified with insect Sf21 cells essentially as described previously (Li et al., 2012). Instead of gel filtration chromatography, buffer was exchanged to 1× MRB80 containing 100 mM KCl and 1 mM DTT, using a desalting column PD MiniTrap G-25 (GE Healthcare). His-GFP-CLAS^Mast/Orbit^ (wild-type and TOG mutant [W334E, K421A, W852E, K940A, which reside in the TOG2 and TOG3 domains]), His-CLASP^Mast/Orbit^, TagRFP-CLASP, and His-Plk1^Polo^(wild-type and K54A kinase-dead mutant) were purified in an identical manner to Sentin, except that the lysis buffer contained 100 mM KCl and 0.1–0.2% Tween-20 (CLASP) or cells were treated with 100 nM okadaic acid 1 h prior to collection (Plk1^Polo^). Purified S2 tubulin, EB1, kinesin-13^Klp10A^, and Plk1^Polo^ were flash frozen (with 20% glycerol for EB1 and 20% sucrose for kinesin-13^Klp10A^ and Plk1^Polo^), whereas other proteins were kept on ice and used within 48 h. Recombinant Cdk1-cyclin B complex was purchased from New England Biolab. Gel filtration chromatography of GFP-CLASP^Mast/Orbit^ was performed with a Superdex 200 increase 10/300 GL column equilibrated with a gel filtration buffer (80 mM K-Pipes, pH 6.8, 100 mM KCl, 4 mM MgCl2, 1 mM EGTA, and 1 mM DTT).

### In vitro MT polymerisation assay

The *in vitro* MT polymerisation assay was performed basically following the method described in (Li et al., 2012), which referred to (Bieling et al., 2010; Gell et al.,2010). Flow chambers (22 mm [width] × 1 mm [height] × 0.15 mm [depth]) were assembled between a coverslip and a precleaned micro slide glass with double-sided tape. Glass was soaked for 3 days in water supplied with the detergent SCAT 20-X, rinsed with fresh water, fired with methanol, and then silanized. The silanized coverslip was coated with anti-biotin (1–5% in 1× MRB80, Invitrogen), and the nonspecific surface was blocked with Pluronic F127 (1% in 1x MRB80, Invitrogen). Biotinylated MT seeds (50–100 μM tubulin mix containing 10% biotinylated pig tubulin and 10% Alexa568-labelled pig tubulin with 1 mM GMPCPP) were specifically attached to the functionalised surface by biotinylated tubulin-anti-biotin links. After the chamber was washed with 1× MRB80, MT growth was initiated by flowing tubulin (containing 3.3% Alexa568-labelled pig tubulin), EB1 (400 nM), Sentin (200 nM), GFP-CLASP^Mast/Orbit^ (0–45 nM), kinesin-13^Klp10A^ (100 nM, with 1 mM ATP), and/or XMAP215^Msps^-HA (100 nM) into the assay buffer (1x MRB80, 75 mM KCl, 1 mM GTP, 0.5 mg/mL k-casein and 0.1% methylcellulose), and an oxygen scavenger system (50 mM glucose, 400 μg/mL glucose-oxidase, 200 μg/mL catalase, and 4 mM DTT).

The samples were sealed with candle wax. We used fixed concentrations of EB1, Sentin, and XMAP215^Msps^ based on our previous study, where the effect of these three factors combined has been demonstrated at these concentrations (Li et al., 2012). Kinesin-13^Klp10A^ was used at 100 nM, since increasing this protein further depolymerised virtually all MT seeds in the imaging field. To match the estimated cellular tubulin concentration (Fig. S5), we used 15 μM S2 tubulin for our main reconstitution experiments involving five or four non-tubulin proteins (Fig. 2 and 3). However, in some experiments, we used 10 μM S2 tubulin to save the protein whose yield was low compared to yields of pig brain-derived tubulin (Fig. 1B and 5B, D). When the samples were treated with Cdk1-cyclinB or Plk1^Polo^ kinase, 5 or 100 nM of recombinant kinase (with 2 mM ATP), respectively, was added to the reaction. During the experiments, the samples were kept at 27 °C; images were collected every 3 s for 20 min via TIRF microscopy with a Ti system (Nikon), EMCCD camera (Evolve; Roper), 100× (1.49 NA) lens, and a 488/561-nm excitation laser. The microscopes were controlled by Micromanager.

### Data analysis

All parameters of MT plus-end dynamics *in vitro*, except those related to pause, were determined in accordance with the method described in (Li et al., 2012). In brief, kymographs were generated for every MT seed identified in the image field, and ∽10 MTs, with a traceable plus end, were randomly selected. The starting and end points of growth and shrinkage were identified in the kymograph. The duration of and change in MT length for each phase were measured, and the rate was determined by calculating the mean velocity between the start and end-point of each phase. Catastrophe frequency was determined by dividing the number of shrinkage events by the sum of growth and pause time. The transition from shrinkage to pause or growth was considered a rescue event, and the rescue frequency (for shrinkage time) was calculated. When MTs did not grow or shrink more than 2 pixels (∽0.35 μm) for ≥5 frames (15 sec), this period was defined as pause; this definition was comparable to that used in the *in vivo* pause measurement of our previous study (Li et al., 2011). In the analysis presented in Fig. 1 and S1, we did not count the pause phase, because it was impossible to identify the starting point of the pause phase of these slowly and persistently growing MTs; for them we obtained the growth rate throughout. Catastrophe frequency in the absence of kinesin-13^Klp10A^ was measured by directly tracing the MT plus ends, since MTs were overly crowded and kymographs could not be generated.

### Data presentation

We followed (Li et al., 2012) for data presentation. As was described in (Li et al., 2012), some degrees of variation in MT behaviours were observed on different experiment days, possibly due to differences in protein batches. To adjust for this variation, we have provided most of the graphs for the values for each condition relative to the control data acquired on the same day. In contrast, the mean values of three experiments have been provided in Tables. For example, when we obtained growth rates of a_1_, a_2_, and a_3_ (μm/min) in three independent experiments using tubulin alone, and b_1_, b_2_, and b_3_ (μm/min) after CLASP^Mast/Orbit^ addition, we presented the relative values as b_1_/a_1_, b_2_/a_2_, and b_3_/a_3_ in grey and (b_1_/a_1_ + b_2_/a_2_ + b_3_/a_3_)/3 in black in the figure. In Tables, in contrast, we provided the mean rates of (a_1_+a_2_+a_3_)/3 and (b_1_+b_2_+b_3_)/3 separately with SDs. For all parameters, the p-values based on a Student’s t-test analysis of 3–4 experiments were obtained and presented in Tables 1 and S1–S3. However, the statistical significance of these differences could not be properly discussed since sample sizes were small; nevertheless, the values served as a good indicator of the degree of change. For growth and shrinkage rates where sufficiently large numbers of MTs were analysed, we additionally provided p-values for each experiment based on a Mann-Whitney U-test (therefore, three or four p-values are displayed in each column).

### Cell biology

S2 cells that express EB1-GFP and mCherry-tubulin were cultured and RNAi was utilised as previously described (Bettencourt-Dias and Goshima, 2009; Goshima et al., 2007). For Klp61F RNAi, cells were treated with dsRNAs for 72 h and then plated on ConA-coated glass-bottom dishes for microscopy (Goshima and Vale, 2003; Goshima et al., 2007). Live imaging of EB1-GFP was performed using spinning-disc confocal microscopy with a 100× 1.40 NA or 100× 1.45 NA objective lens (∽25 °C). A CSU-X1 confocal unit (Yokogawa, Japan) attached to a Nikon TE or Ti inverted microscope, and EMCCD camera ImagEM (Hamamatsu, Japan) were used for image acquisition. The microscopes and attached devices were controlled using Micromanager. To eliminate actin cytoskeleton that might have physically prevented MT growth (Li et al., 2011), an actin inhibitor (latrunculin A, 2.5 μM) was added to cells prior to imaging. In the Plk1^Polo^ inhibition experiment, cells were treated with control DMSO or 1 μM BI 2536, which was shown to be effective in S2 cells (Januschke et al., 2013; Kachaner et al., 2014), and images were acquired at 3-s intervals.

### Phos-tag SDS-PAGE, autoradiography, and immunoblotting

Phosphorylation of purified proteins by Plk1^Polo^, except CLASP^Mast/Orbit^, was evaluated by Phos-tag SDS-PAGE (Kinoshita et al., 2006). 1 mM ATP was added to the reaction. EB1 was run on an 8% acrylamide gel, whereas a 5% gel was used for tubulin, Sentin, and kinesin-13^Klp10A^. The gel for XMAP215^Msps^ was a mixture of 3% acrylamide and 0.5% agarose, wherein agarose assisted in gel polymerisation. Acrylamide conjugated with the Phos-tag molecule at 20 μM. For unknown reasons, the GFP-CLASP^Mast/Orbit^ band was not clearly detectable on any type of gel. CLASP^Mast/Orbit^ phosphorylation was therefore monitored by autoradiography, where 113 nM Plk1^Polo^ (wild-type or KD) was mixed with 1 μM GFP-CLASP^Mast/Orbit^ in MRB80/DTT with 100 mM KCl, 0.1 mM ATP (containing 110 nM ATP-P^32^), and 80 mM β-glycerophosphate for 30 min at 24–25 °C, followed by SDS-PAGE and Coomassie staining. Autoradiography was used to test for phosphorylation by Cdk1, where 27 nM recombinant Cdk1-cyclin-B was mixed with 2.1 μM EB1, 1.6 μM tubulin, 1.7 μM Sentin, 1.8 μM kinesin-13^Klp10A^, 1.2 μM CLASP^Mast/Orbit^, or 2.2 μM XMAP215^Msps^. Immunoblotting of the S2 extract was performed with the following antibodies, anti-α-tubulin (DM1A; Sigma; 1:1000), anti-CLASP^Mast/Orbit^ (1:300; gift from Dr. Helder Maiato [University of Porto]), anti-EB1 (1:1000; gift from Dr. Stephen Rogers), anti-Klp10A (1:300; gift from Dr. David Sharp), anti-Sentin (1:1000; (Li et al., 2011)), and anti-XMAP215^Msps^ (1:500; gift from Dr. Hiroyuki Ohkura).

## Acknowledgements

We wish to thank Tomomi Tani (Marine Biological Laboratory, Woods Hole, MA), Wenjing Li (Tsinghua University, China), Momoko Nishina, Rie Inaba, Tomoya Edzuka, and Tomoko Nishiyama (Nagoya University, Japan) for technical assistance, and Wenjing Li and Tomomi Kiyomitsu (Nagoya University) for valuable comments regarding the manuscript. This work was supported by the Grants-in-Aid for Scientific Research (KAKENHI; 15H01317, 15KT0077). T.M. is a recipient of a JSPS pre-doctoral fellowship.

---

**Table S1.** Kinetic parameters of *in vitro* reconstituted MT polymerisation dynamics with or without Sentin (graphic presentation in Fig. 3)

| Condition | N | Growth rate | Shrinkage rate | Catastrophe frequency | Rescue frequency |
| --- | --- | --- | --- | --- | --- |
| | | ($\mu\text{m}/\text{min}$) | ($\mu\text{m}/\text{min}$) | ($\times 10^{-3}\text{s}^{-1}$) | ($\times 10^{-3}\text{s}^{-1}$) |
| +Sentin | 3 | 4.7 ± 0.8 (283) | 8.0 ± 1.7 (239) | 24.0 ± 10.2 (227) | 85.7 ± 32.9 (232) |
| -Sentin | 3 | 2.0 ± 0.4 (179) | 6.1 ± 1.5 (92) | 4.4 ± 2.3 (85) | 95.0 ± 26.1 (86) |
| P-values |  | 0.006 (n = 3)<br><10<sup>-4</sup> (exp1)<br><10<sup>-4</sup> (exp2)<br><10<sup>-4</sup> (exp3) | 0.227 (n = 3)<br>0.062 (exp1)<br>0.001 (exp2)<br>0.040 (exp3) | 0.032 (n = 3) | 0.719 (n = 3) |

Mean ± SD of three independent experiments (total number of events).

**Table S2.** Kinetic parameters of *in vitro* reconstituted MT polymerisation dynamics with or without Cdk1 (graphic presentation in Fig. S4)

| Condition | N | Growth rate | Shrinkage rate | Catastrophe frequency | Rescue frequency |
| --- | --- | --- | --- | --- | --- |
| | | ($\mu\text{m}/\text{min}$) | ($\mu\text{m}/\text{min}$) | ($\times 10^{-3}\text{s}^{-1}$) | ($\times 10^{-3}\text{s}^{-1}$) |
| -Cdk1 | 3 | 4.7 ± 2.0 (181) | 4.9 ± 0.7 (135) | 9.7 ± 4.6 (120) | 74.8 ± 18.4 (135) |
| +Cdk1 | 4 | 5.0 ± 1.8 (251) | 6.2 ± 1.1 (184) | 12.4 ± 6.9 (161) | 57.8 ± 17.6 (184) |
| P-values |  | 0.73 (n = 3, 4)<br><10<sup>-4</sup> (exp1)<br>0.085* (exp2)<br>0.011 (exp3)<br>0.102 (exp4) | 0.13 (n = 3, 4)<br>0.307# (exp1)<br>0.007 (exp2)<br>0.625 (exp3)<br>0.001 (exp4) | 0.63 (n = 3, 4) | 0.38 (n = 3, 4) |

Mean ± SD of three independent experiments (total number of events). \*In this experiment, the growth rate was decreased by Cdk1 treatment. #In this experiment, the shrinkage rate was decreased by Cdk1 treatment

**Figure. S1.**
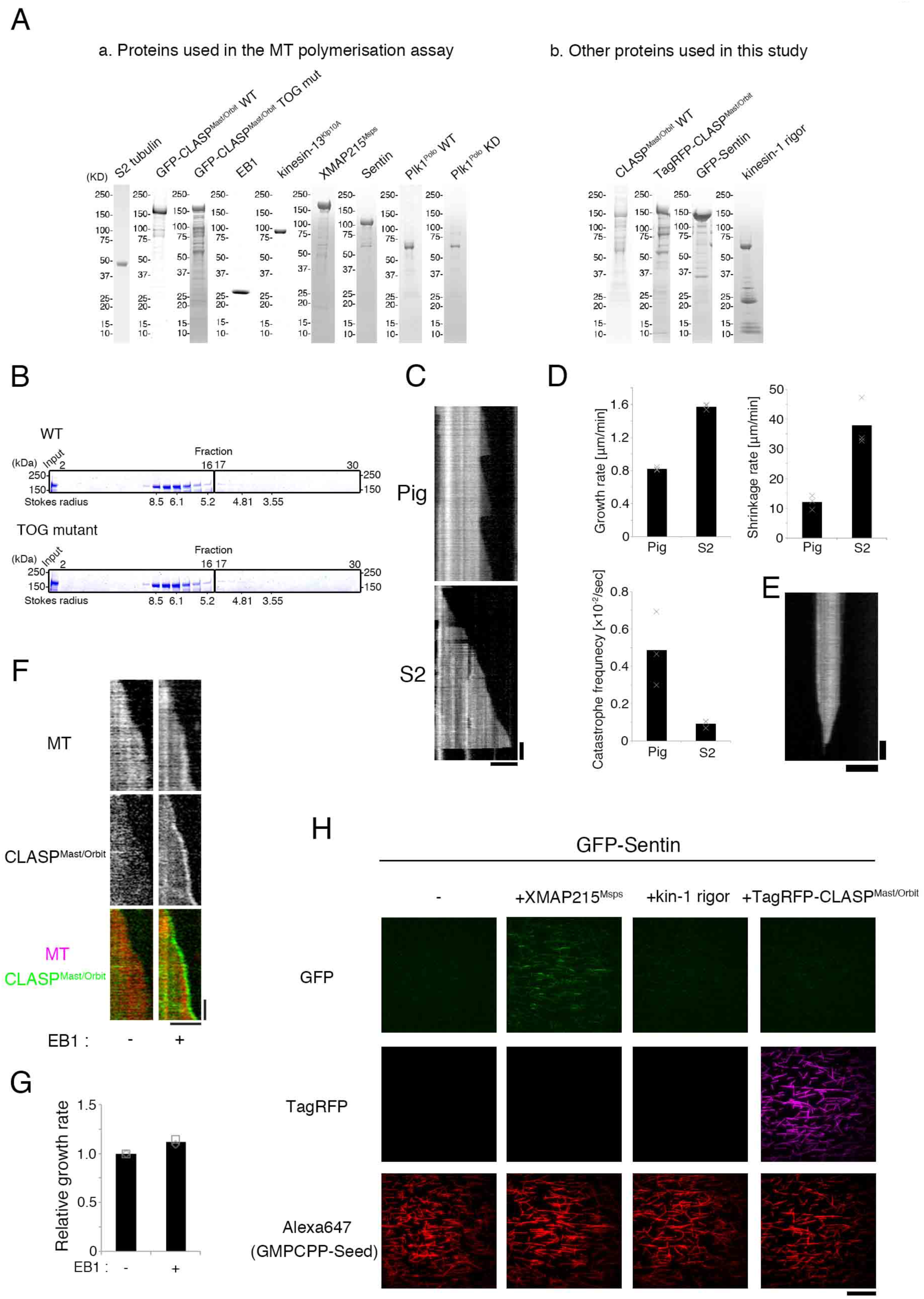
Recombinant proteins used in this study, kinesin-13^Klp10A^ activity, and comparison between S2-derived tubulin and pig brain tubulin. (**A**) Coomassie staining of recombinant proteins used in this study. **(B)** Gel filtration chromatography of GFP-CLASP^Mast/0rbit^ (wild-type and TOG mutant) followed by Coomassie staining. Molecular size markers (thyloglobulin, ferritin, catalase, aldolase, BSA) were run to estimate the Stokes radius. (**C**) Kymographs showing MT polymerisation and depolymerisation with pig or S2 tubulin. **(D)** Parameters of MT polymerisation dynamics in the presence of 15 μM S2- or pig tubulin alone. The mean values of each experiment are marked in grey, whereas the mean values of all the experiments are indicated by black graphs. Actual values are plotted in these graphs. Pausing and rescue were rarely observed in either condition. **(E)** Kymograph showing that kinesin-13^Klp10A^ depolymerises MT seeds. 100 nM kinesin-13^Klp10A^ was mixed with GMPCPP-MT seeds and 10 μM taxol. Horizontal bars 5 μ Vertical bars, 1 min. **(F)** Kymographs showing EB1-dependent tip accumulation of GFP-CLASP^M 0rblt^. 400 nM EB1 and 100 nM CLASP^Mast/01tlt^ were mixed with 30 μM pig tubulin. Horizontal bar, 5 μm; Vertical bar, 1 min. (**G**) Comparison of the MT growth rate in the presence or absence of EB1. Catastrophe was almost completely suppressed in either condition. **(H)** CLASP^Mast/Orbit^ does not recruit Sentin to the MT lattice. TagRFP-CLASP^Mast/Orbit^ (45 nM) was mixed with GFP-Sentin (10 nM) and MT seeds. Although M act/Or hit TagRFP-CLASP^Mast/Orbit^localised to the MT lattice, as expected, GFP-Sentin signals were invisible. An unrelated MT-associated protein (kinesin-1 rigor mutant [560 a.a.] (Rice et al., 1999)) and XMAP215^Msps^, which directly binds to Sentin (Li et al., 2012), were used as negative and positive controls, respectively (each 100 nM). Bar, 20 μm.

**Figure. S2.**
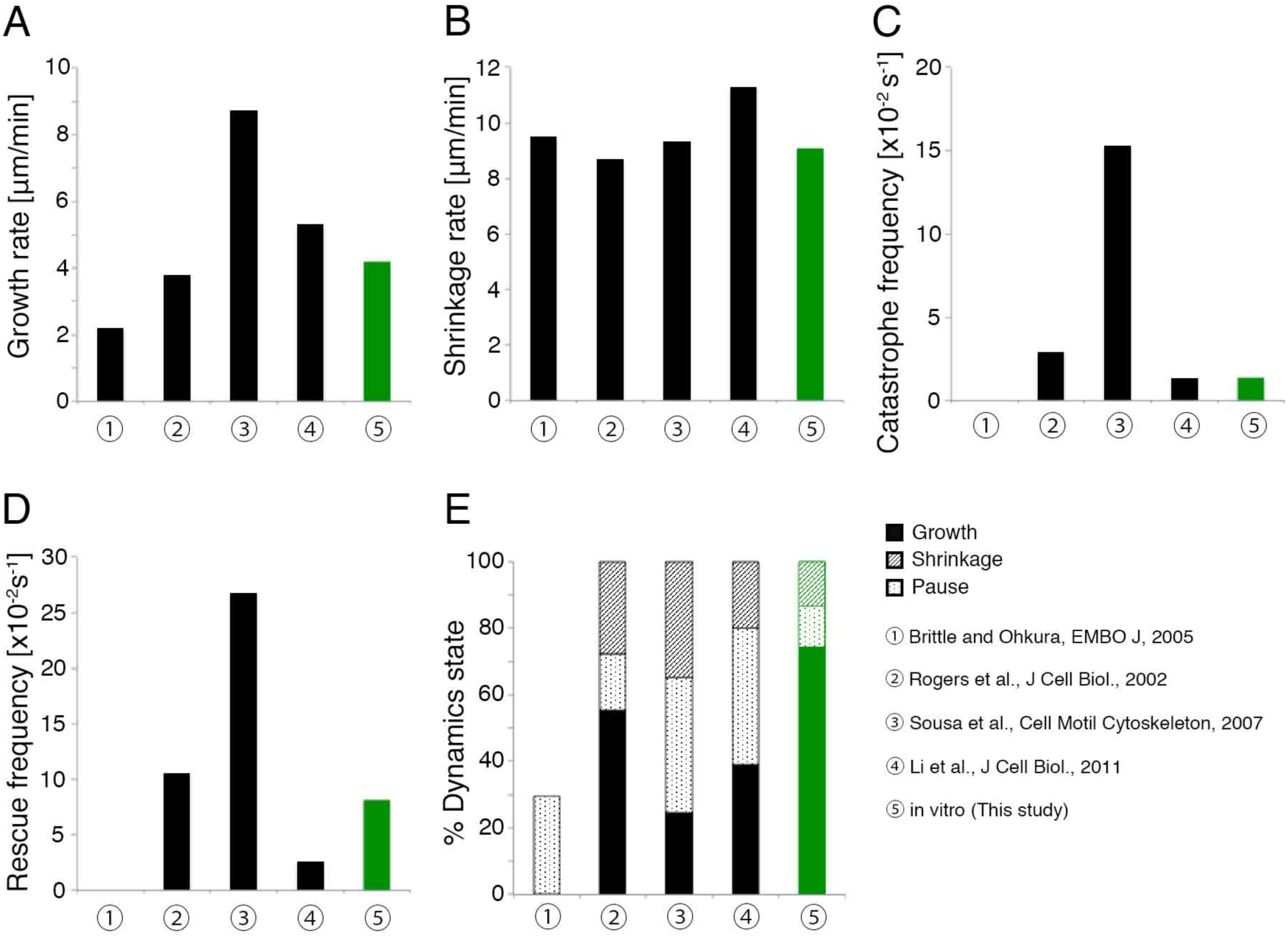
Comparison of *in vitro* and *in vivo* parameters associated with MT dynamics. Kinetic parameters of S2 interphase cytoplasmic MT dynamics presented in the previous four studies and our current *in vitro* study (data from Fig. 2, 15 nM CLASP^Mast/Orbit^). Note that catastrophe or rescue frequency was obtained by dividing event numbers by total time in the *in vivo* studies. We recalculated this parameter based on the formula used in this study; event numbers were divided by total growth/pause or shrinkage time.

**Figure. S3.**
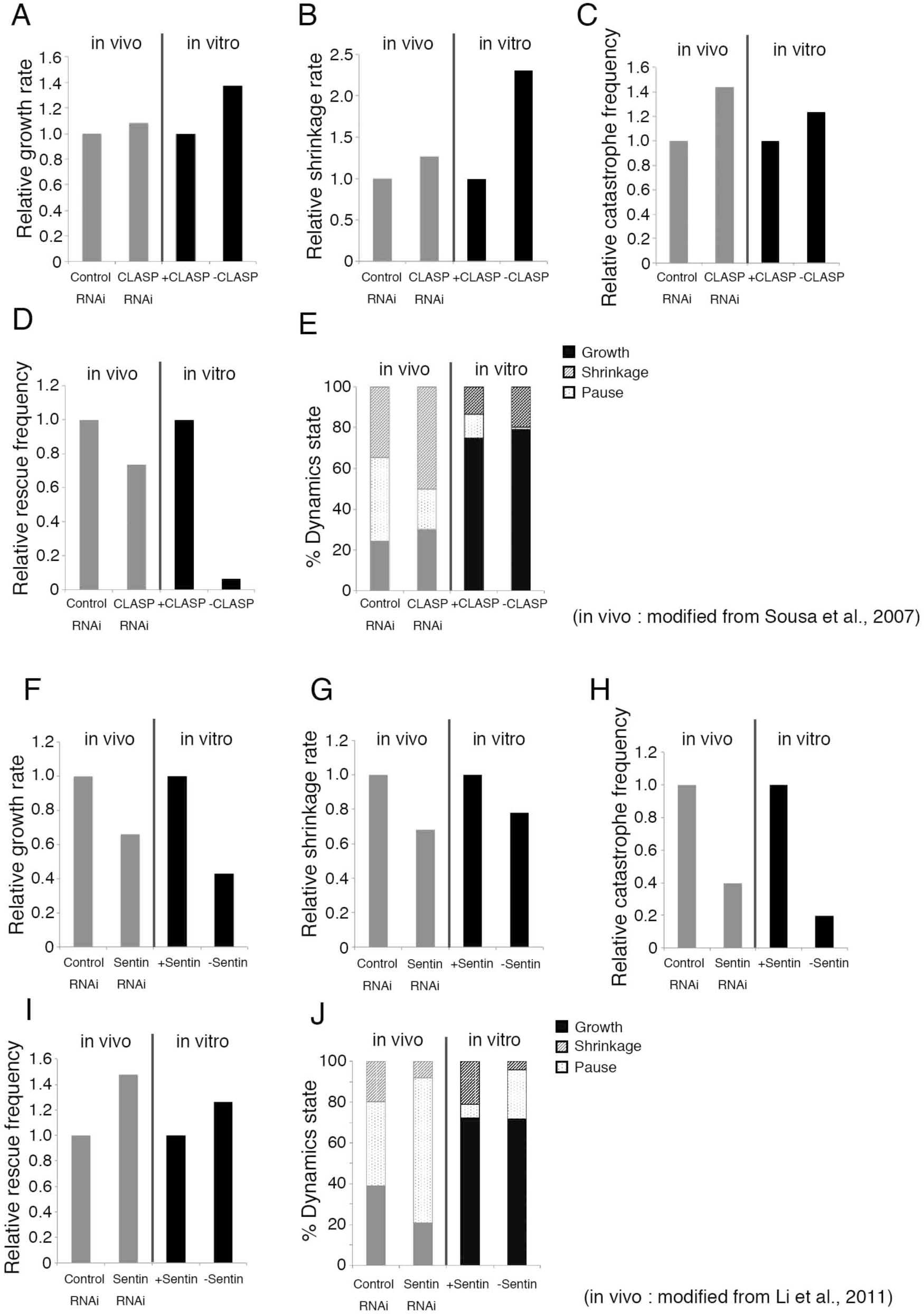
Comparison of the *in vitro* and *in vivo* effects of CLASP^Mast,0rbit^ and Sentin. (**A-E**) Comparison of the change in kinetic parameters obtained *in vivo* (control vs.CLASP^Mast/0rbit^ RNAi(72 h); (Sousa et al., 2007)) and *in vitro* (five factors vs. CLASP^Mast/0rbit^-depleted four factors). (**F-J**) Comparison of the change in kinetic parameters obtained *in vivo* (control vs. Sentin RNAi; (Li et al., 2011)) and *in vitro* (five factors vs. Sentin-depleted four factors). Note that catastrophe or rescue frequency was obtained by dividing event numbers by total time in the *in vivo* studies. We re-calculated this parameter based on the formula used in this study; event numbers were divided by total growth/pause or shrinkage time.

**Figure. S4.**
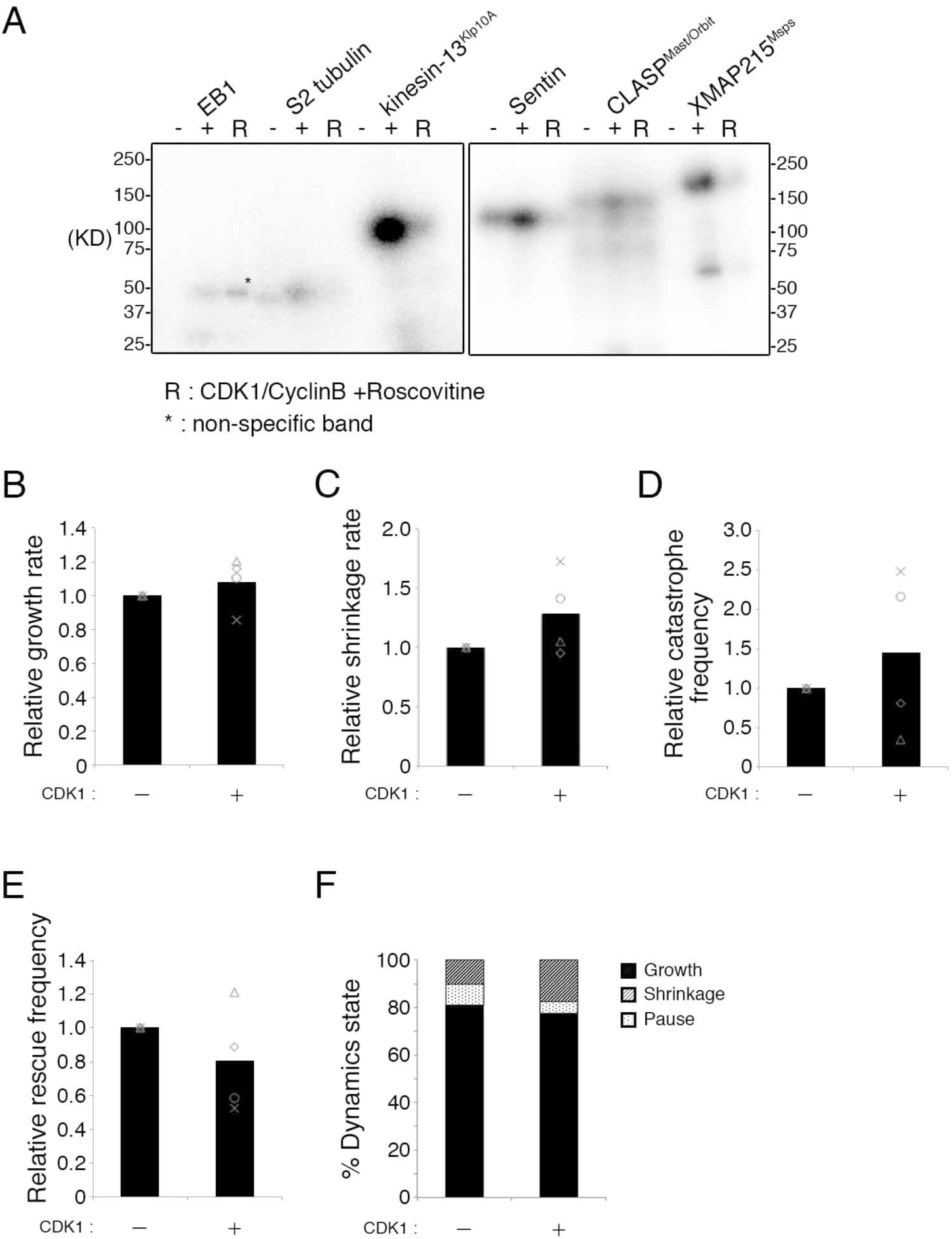
Effect of Cdk1-cyclinB in MT dynamics *in vitro*. (**A**) Autoradiography indicated that the Cdk1-cyclinB complex efficiently phosphorylates XMAP215^Msps^ kinesin-13^Klp10A^ (4.4- and 16-fold increases in band intensity, respectively). Asterisk indicates phosphorylation of an unknown protein that represents contamination after EB1 purification (EB1 is a ∼30 kD protein [Fig. S1A]). (**B-F**) Effect of Cdkl-cyclinB in MT dynamics reconstituted with five factors. Parameters of MT polymerisation dynamics with 15 (cross) or 10 μM (others) S2 tubulin, XMAP215^Msps^, EB1, Sentin, 15 nM CLASP^Mast/0rbit^, kinesin-13^Klp10A^, and 5 nM Cdk1-cyclinB. Relative values are presented in this figure, whereas the actual values are described in Table S2. The mean values of each experiment are marked in grey, whereas the mean values of all the experiments are indicated by black graphs. In all four experiments, the ‘Cdk−’ condition was created by adding 50 μM roscovitine (Cdk1 inhibitor) simultaneously with recombinant Cdk1 (inhibition of the kinase activity at this roscovitine concentration hadbeen confirmed). In two experiments (marked by a cross and triangle), the ‘Cdk+’ condition was created by DMSO addition instead of roscovitine. In other two experiments (marked by a circle and diamond), the ‘Cdk+’ condition was created by first incubating the protein mixture with Cdk1 for 15 min, followed byroscovitine addition. Note that MTs were more prone to polymerisation in this experiment, since glycerol concentration in the assay buffer was higher (Cdk1 was stored in the buffer containing glycerol). Therefore, catastrophe was a rare event compared to other experiments; the difference in catastrophe frequency in the presence or absence of roscovitine should therefore be interpreted with caution.

**Figure. S5.**
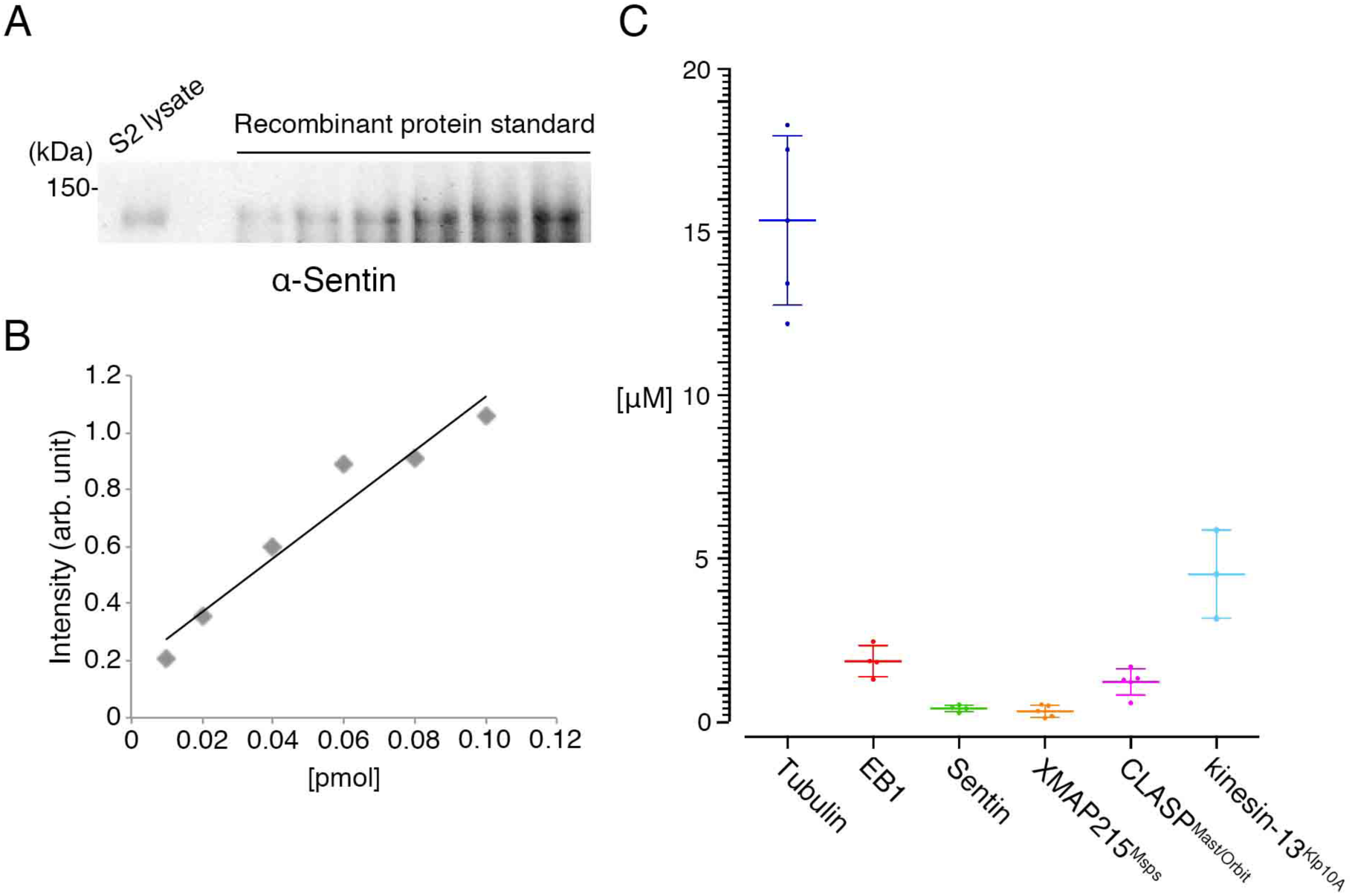
Estimate of ceiiular protein concentration. (**A,B**) An example of quantitative immunoblotting. In this case, the total cell lysate of 2 × 10^4^ S2 cells and 5–50 fmol recombinant His-Sentin were loaded and immunoblotting was performed with an anti-Sentin polyclonal antibody. Intensity of each band corresponding to Sentin protein was measured and plotted. (**C**) Rough estimates of cellular protein concentration using the approximate volume of the S2 cytoplasm (spherical nuclear volume [7.5 μm diameter] was subtracted from spherical S2 cell volume [11.8 μm diameter]). Experiments were performed two to three times, and immunoblotting and band intensity measurements were duplicated (each point in the graph shows a result of each measurement). Mean and SD values are presented.

